# Transcriptional feedback targeting Wnt pathway components reveals hidden heterogeneity in *C. elegans* seam cell lineages

**DOI:** 10.64898/2026.03.04.709659

**Authors:** Mar Ferrando-Marco, Simon Berger, Michalis Barkoulas

## Abstract

Asymmetric cell division in the epidermal stem cells of *Caenorhabditis elegans,* known as seam cells, relies on the Wnt/β-catenin asymmetry pathway to generate daughter cells with distinct fates. However, whether components of this pathway components are transcriptionally regulated during these divisions remains unclear. Here, we employ single molecule fluorescence in situ hybridisation to quantify mRNA distributions of key Wnt pathway components during L2 symmetric and asymmetric seam cell divisions. We find that transcripts encoding the negative regulators *pry-1*/Axin and *apr-1*/APC are enriched in posterior daughter cells, while those encoding the positive regulators *sys-1*/β-catenin, *wrm-1*/β-catenin, and *lit-1*/NLK, along with the transcription factor *pop-1*/TCF, are enriched in anterior daughter cells. Strikingly, molecular asymmetries are already evident following the L2 symmetric division, with anterior and posterior daughters exhibiting distinct levels of Wnt component expression and Wnt pathway activation. These mRNA distributions are surprising considering the established protein localisations that underpin the Wnt asymmetry model and suggest extensive post-divisional transcriptional regulation. We further demonstrate that *pop-1* asymmetric expression depends on Wnt signalling activity, supporting a model in which transcriptional feedback reinforces cell fate decisions. Investigation of protein distributions using knock-in reporters for PRY-1 and CAM-1 showed that protein accumulation patterns at L2 are consistent with transcript levels. Our findings uncover pervasive transcriptional feedback within the Wnt pathway that likely contributes to robust fate specification and reveal molecular heterogeneity with potential functional consequences for lineage behaviour.

## Introduction

The Wnt signalling pathway is an evolutionarily conserved cascade that controls cell polarity and fate decisions in the context of asymmetric cell division across diverse organisms (Bielen & Houart, 2014; Habib & Acebrón, 2022; Lam & Phillips, 2017). Asymmetric cell division is a fundamental mechanism by which stem and progenitor cells generate cellular diversity, producing one daughter that maintains stem/progenitor fate and one that differentiates. Maintaining an appropriate balance between self-renewal and differentiation is essential for tissue homeostasis (Knoblich, 2008; Morrison & Kimble, 2006). In *Caenorhabditis elegans*, the lateral epidermal seam cells provide a powerful model for investigating Wnt-dependent control of cell polarity and fate determination. These cells are arranged in two bilateral rows running along the anterior-posterior axis and undergo a stereotypical series of symmetric and asymmetric divisions during larval development, generating most of the syncytial epidermis and certain neuronal precursor cells (Figure 1A) (Chisholm & Hsiao, 2012; Joshi et al., 2010; Sulston & Horvitz, 1977). During asymmetric divisions, the posterior daughter typically retains seam cell identity and continues dividing in subsequent larval stages, whereas the anterior daughter exits the stem cell-like programme, differentiates and fuses with the surrounding hyp7 syncytium. A single symmetric division in selected seam cell lineages at the L2 stage expands the seam cell population from ten to sixteen per lateral side, a number that is highly consistent in wild-type populations (Boukhibar & Barkoulas, 2016; Katsanos et al., 2017; Koneru et al., 2021). Following the final larval stage, seam cells undergo terminal differentiation, fuse into a syncytium, and secrete the cuticular ridges known as alae.

**Figure 1.**
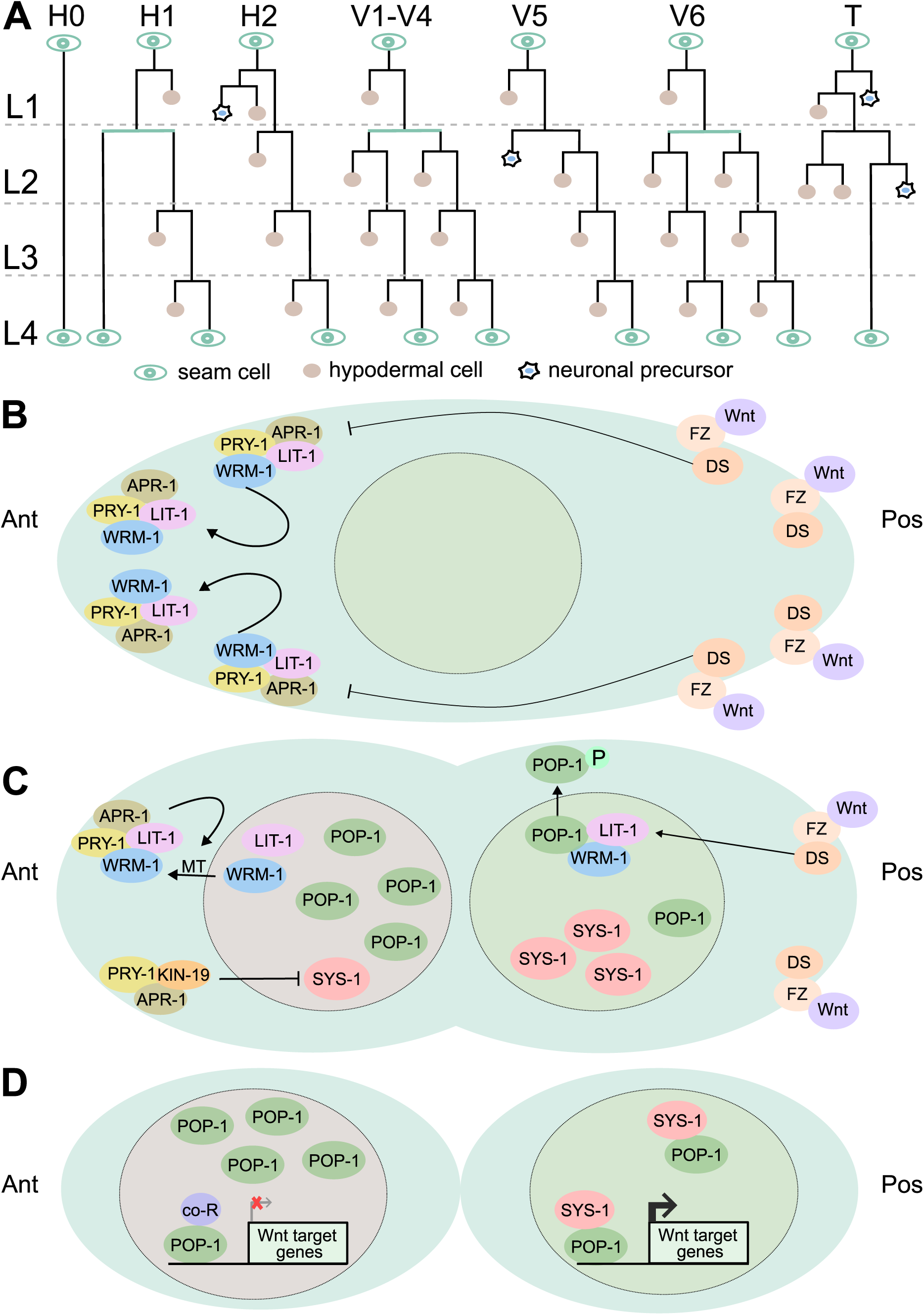
The Wnt/β-catenin asymmetry pathway controls the polarity of asymmetric seam cell divisions. **(A)** Schematic showing seam cell divisions from L1 to L4. Seam cells are labelled in green, hypodermal cells are labelled in brown and neuroblasts are labelled in blue. **(B)** In the mother cell, positive regulators Frizzled (FZ) and Dishevelled (DS) localise to the posterior cortex upon Wnt signal reception, while negative regulators APR-1/APC and PRY-1/Axin localise to the anterior cortex, recruiting WRM-1/β-catenin and LIT-1/NLK. Cortical WRM-1 further recruits APR-1 and WRM-1. **(C)** In the dividing cell, cortical APR-1 stabilises microtubules (MT), facilitating the export of WRM-1 and LIT-1 from the anterior nucleus. APR-1, PRY-1, and KIN-19/CKI promote SYS-1/β-catenin degradation in anterior daughter cells, resulting in asymmetric localisation towards the posterior nucleus. WRM-1 and LIT-1 phosphorylate POP- 1/TCF in the posterior nucleus, triggering its nuclear export, with LIT-1 activity and localisation dependent on MOM-4/MAPK and WRM-1. **(D)** In anterior daughter cells, high nuclear POP-1 recruits co-repressors (co-R) to repress Wnt target genes. In posterior daughter cells, low nuclear POP-1 forms a complex with SYS-1 to activate Wnt target gene expression.

The Wnt/β-catenin asymmetry pathway plays a central role in this process by globally regulating the fate of seam cell daughters: reduced Wnt signalling causes both daughters to adopt differentiated fates, while constitutive pathway activation leads to symmetric retention of the seam cell fate and supernumerary seam cells (Gleason & Eisenmann, 2010; Gorrepati et al., 2013). In this modified version of the Wnt pathway, Wnt component localisation and protein stability are tightly regulated to generate asymmetric protein distribution between daughter cells (Mizumoto & Sawa, 2007b, 2007a; Phillips et al., 2007; Takeshita & Sawa, 2005), leading to differentiation of the anterior daughter cell and retention of seam cell fate in the posterior daughter cell. In the dividing mother cell, Wnt ligands promote the posterior cortical localisation of Frizzled and Ror receptors together with Dishevelled proteins, while the negative regulators APR-1/APC and PRY-1/Axin localise to the anterior cortex (Figure 1B) (Baldwin & Phillips, 2014; Goldstein et al., 2006; Green et al., 2008; Mizumoto & Sawa, 2007a). At the anterior cortex, APR-1 recruits WRM-1/β-catenin, which in turn recruits the kinase LIT-1/NLK, establishing a feed-forward loop that reinforces cortical asymmetry (Figure 1B) (Mizumoto & Sawa, 2007b, 2007a; Sugioka et al., 2011; Takeshita & Sawa, 2005). During division, APR-1 also promotes the export of WRM-1 and LIT-1 from the anterior nucleus through regulation of microtubule dynamics and, together with KIN-19/CKIα, promotes degradation of SYS-1/β-catenin (Figure 1C) (Baldwin & Phillips, 2014; Banerjee et al., 2010; Mizumoto & Sawa, 2007a; Sugioka et al., 2011). In posterior daughter cells, the absence of anterior negative regulators allows stabilisation of SYS-1 and nuclear retention of WRM-1 and LIT-1. High nuclear WRM-1 promotes activation of LIT-1 leading to phosphorylation of the TCF transcription factor POP-1 and its nuclear export (Figure 1C) (Lo et al., 2004; Meneghini et al., 1999; Takeshita & Sawa, 2005; Yang et al., 2011, 2015). The resulting ratio of nuclear levels of POP-1 and SYS-1 is thought to determine transcriptional output in the daughter cells, with low POP-1 and high SYS-1 in posterior daughters promoting transcriptional activation, and high POP-1 and low SYS-1 in anterior daughters favouring transcriptional repression (Figure 1D) (Bekas & Philips, 2022; Huang et al., 2007; Phillips et al., 2007).

To date, the Wnt/β-catenin asymmetry pathway has been characterised primarily through analyses of protein localisation through translational reporters imaged at early stages. Therefore, to what extent components of this pathway are also regulated transcriptionally during division remains unclear. Here, we use single molecule fluorescence in situ hybridisation (smFISH) to provide a comprehensive transcriptional analysis of key Wnt/β-catenin asymmetry pathway components in V1-V4 seam cell lineages, which show identical division patterns. We focus on the L2 stage to capture both symmetric and asymmetric division events. We uncover unexpected asymmetric mRNA expression patterns following division, with the negative regulators *pry-1* and *apr-1* enriched in posterior seam-fated daughters and the positive regulators *sys-1*, *wrm-1* and *lit-1*, together with the transcription factor *pop-1*, enriched in anterior differentiating daughters. Notably, we find that daughter generated from seemingly symmetric divisions differ significantly in the distributions of Wnt components and in their capacity for Wnt pathway activation, indicating that they are not truly identical and harbour substantial molecular heterogeneity. These transcriptional patterns were largely not anticipated based on the known protein distributions and we demonstrate that they are likely to reflect transcriptional feedback in the pathway. We propose that transcriptional feedback may contribute to robust seam cell fate determination and influence how cells within a seam cell lineage respond to genetic or environmental changes.

## Results

### Negative regulators *pry-1* and *apr-1* are enriched in posterior daughter cells

To characterise the expression pattern of Wnt/β-catenin asymmetry pathway components in seam cells, we performed smFISH focusing on V1-V4 seam cells at late L1 and L2 stages. We first examined the expression patterns of negative regulators *pry-1* and *apr-1* (Figure 2). Analysis of *pry-1* mRNA distribution at late L1 revealed expression in seam cells (p) prior to the L2 symmetric division (Figure 2A, B). Interestingly, following the L2 symmetric division, posterior (pp) cells displayed significantly higher *pry-1* mRNA levels compared to anterior (pa) cells, which is indicative of repression occurring in anterior cells (Figure 2B). The same pattern was observed following the L2 asymmetric division, where posterior daughter cells (pap and ppp) exhibited significantly higher *pry-1* mRNA levels compared to their corresponding anterior sister cells (paa and ppa) and at comparable levels to those observed at late L1 (Figure 2B).

**Figure 2.**
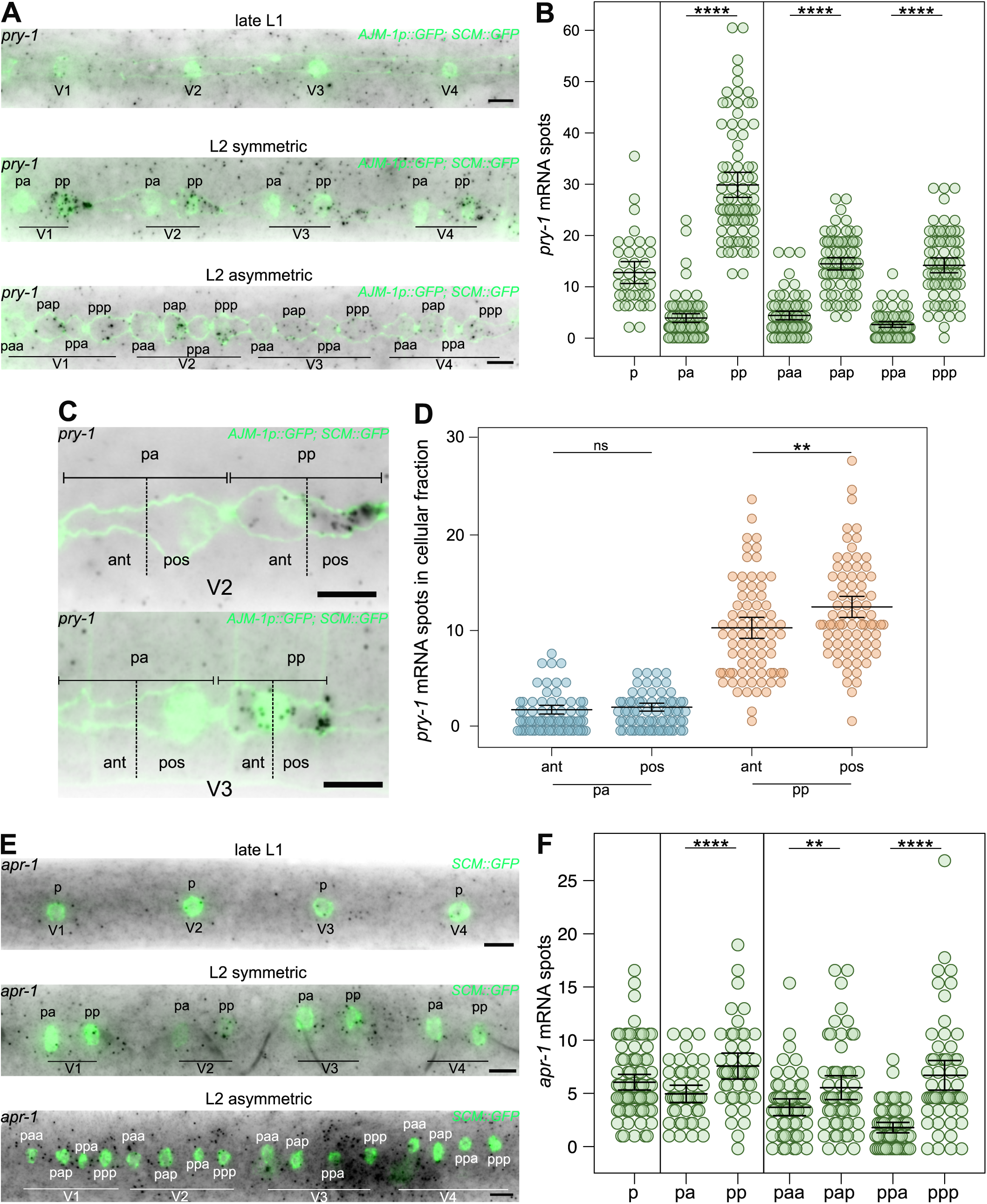
Negative regulators *pry-1* and *apr-1* are enriched in posterior daughter cells. **(A)** Representative *pry-1* smFISH images at late L1, and following the L2 symmetric and L2 asymmetric divisions. **(B)** Quantification of *pry-1* mRNA spots in V1-V4 seam cells before (p) and after the L2 symmetric division (pa, pp), and following the L2 asymmetric division (paa, pap, ppa, ppp); 40 ≤ n ≤ 76 cells per condition. **(C)** Detailed image of *pry-1* smFISH following the L2 symmetric division showing enrichment of mRNAs in the posterior fraction of the cell (pos) compared to the anterior fraction (ant). Anterior and posterior fractions correspond to half of the total length of the cell. **(D)** Quantification of *pry-1* smFISH spots in the anterior and posterior fractions of pa and pp cells following the L2 symmetric division. n = 72 pa cells and 79 pp cells. **(E)** Representative *apr-1* smFISH images at late L1, and following the L2 symmetric and L2 asymmetric divisions. **(F)** Quantification of *apr-1* mRNA spots in V1-V4 seam cells before (p) and after the L2 symmetric division (pa, pp), and following the L2 asymmetric division (paa, pap, ppa, ppp); 44 ≤ n ≤ 80 cells per condition. In A, C and E seam cell nuclei are labelled using *SCM::GFP* and in A and C seam cell membrane with the apical junction marker *ajm-1p::ajm-1::GFP.* Scale bars are 5μm in A, C and E. Error bars in B, D and F show the mean ± standard deviation and **** represent *p*<0.001 and ** *p*<0.01 with a two-tailed t-test.

To further characterise the cause of this asymmetry, we quantified *pry-1* mRNA distribution within individual cells and found that *pry-1* transcripts were enriched in the posterior fraction of pp cells relative to their anterior fraction (Figure 2C, D). This suggests that in the case of *pry-1,* asymmetric localisation of mRNAs may contribute to asymmetric inheritance in daughter cells. Similarly, *apr-1* mRNA expression was enriched in posterior daughter cells following both L2 symmetric and asymmetric divisions (Figure 2E, F) and expression levels in posterior daughter cells following the L2 asymmetric division were like those at late L1 (Figure 2F). Taken together, these results suggest enrichment of negative regulators of Wnt signalling in the posterior seam cells likely due to repression of expression in anterior cells.

### Positive regulators *sys-1*, *wrm-1* and *lit-1* are enriched in anterior daughter cells

We next examined the expression pattern of positive regulators of the Wnt β-catenin asymmetry pathway, namely the β-catenins *sys-1* and *wrm-1*, and the kinase *lit-1*. *sys-1* smFISH analysis revealed expression in seam cells at late L1 (Figure 3A, B). Following the L2 symmetric division, *sys-1* mRNA levels remained comparable between pa and pp cells (Figure 3A, B), whereas following the L2 asymmetric division, anterior daughters (paa and ppa) showed significantly higher *sys-1* mRNA compared to their posterior sisters (pap and ppp), with overall expression levels continuing to decrease relative to earlier stages (Figure 3B). *wrm-1* mRNA levels were significantly higher in anterior daughters compared to posterior daughters following both symmetric and asymmetric divisions pointing to repression in posterior daughter cells (Figure 3C, D). Following the L2 asymmetric cell division, posterior daughter cells presented a strong reduction in *wrm-1* transcript abundance, with 32% of cells showing no *wrm-1* mRNAs at all (Figure 3D). Similarly, to *sys-1*, *wrm-1* mRNA levels were lower at late L2 stage compared to late L1 (Figure 3D). In contrast to *sys-1* and *wrm-1*, *lit-1* showed similar expression across L1 and L2 stages and following the L2 symmetric and L2 asymmetric divisions *lit-1* showed a mild enrichment in anterior daughter cells (Figure 3E, F). Taken together, these results suggest that positive regulators, with *wrm-1* being the most striking example, show enrichment in anterior cells following division likely due to repression in posterior cells.

**Figure 3.**
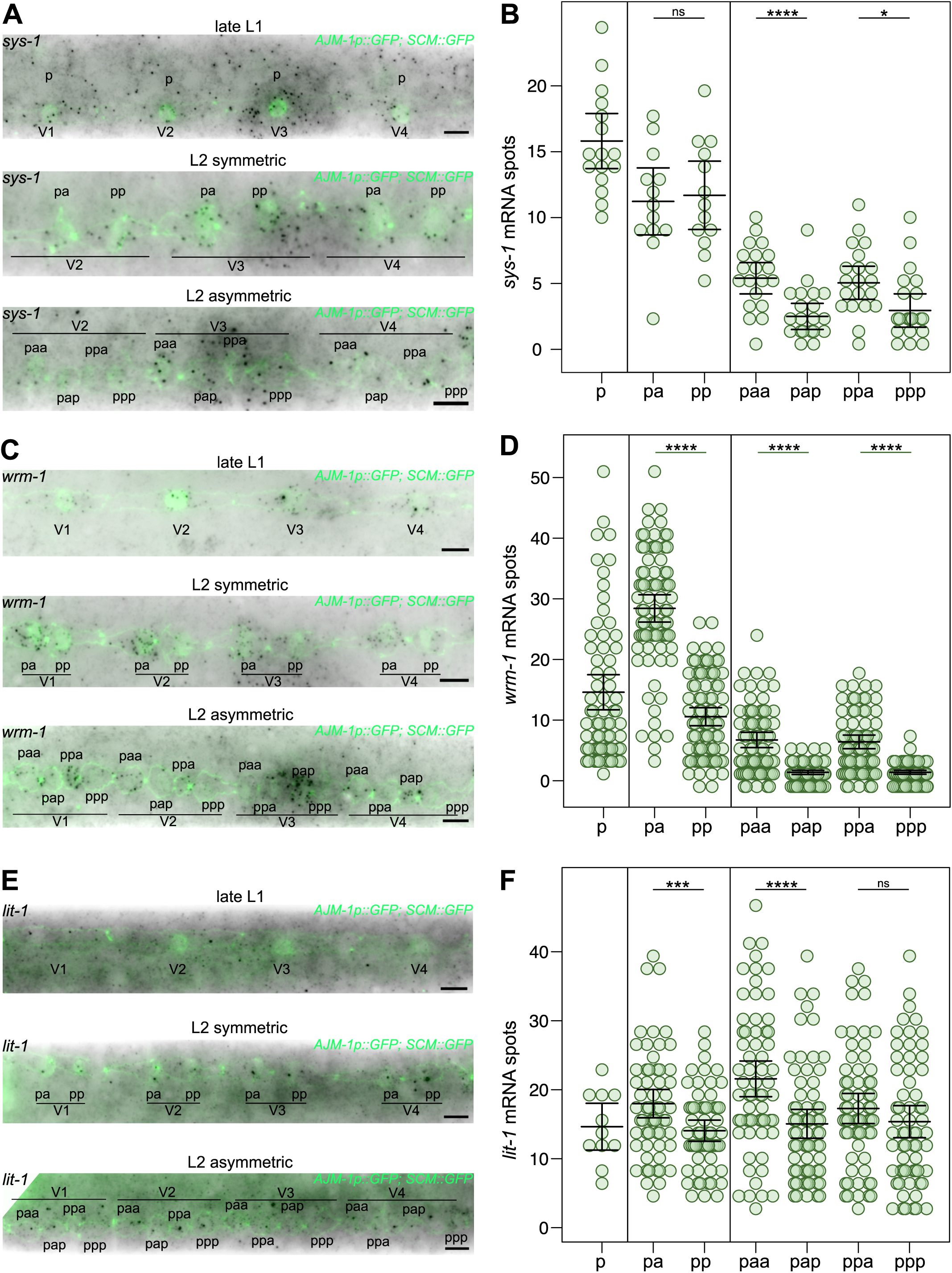
Positive regulators *sys-1*, *wrm-1* and *lit-1* are enriched in anterior daughter cells. **(A)** Representative *sys-1* smFISH images at late L1, and following the L2 symmetric and L2 asymmetric divisions. **(B)** Quantification of *sys-1* mRNA spots in V1-V4 seam cells before (p) and after the L2 symmetric division (pa, pp), and following the L2 asymmetric division (paa, pap, ppa, ppp); 13 ≤ n ≤ 40 cells per condition. **(C)** Representative *wrm-1* smFISH images at late L1, and following the L2 symmetric and L2 asymmetric divisions. **(D)** Quantification of *wrm-1* mRNA spots in V1-V4 seam cells before (p) and after the L2 symmetric division (pa, pp), and following the L2 asymmetric division (paa, pap, ppa, ppp); 67 ≤ n ≤ 78 cells per condition. **(E)** Representative *lit-1* smFISH images at late L1, and following the L2 symmetric and L2 asymmetric divisions. **(F)** Quantification of *lit-1* mRNA spots in V1-V4 seam cells before (p) and after the L2 symmetric division (pa, pp), and following the L2 asymmetric division (paa, pap, ppa, ppp); 11 ≤ n ≤ 63 cells per condition. In A, C and E, seam cell nuclei are labelled using *SCM::GFP* and seam cell membrane with the apical junction marker *ajm-1p::ajm-1::GFP* and scale bars are 5μm. Error bars in B, D and F show the mean ± standard deviation and **** represent *p*<0.001, *** *p*<0.005, and * *p*<0.05 with a two-tailed t-test.

### *pop-1* expression is enriched in anterior daughter cells

These unexpected patterns in the localisation of Wnt machinery suggested a high degree of transcriptional regulation and led us to investigate in more detail the mRNA pattern of the transcription factor POP-1. POP-1 shows a well-described anterior nuclear enrichment at the protein level in daughter cells following asymmetric division (Phillips & Kimble, 2009), as well as a newly reported anterior enrichment at the transcript level (Ferrando-Marco & Barkoulas, 2025). At late L1, seam cells showed *pop-1* mRNA expression that served as the baseline for temporal comparisons (Figure 4A-i, B). During the L2 symmetric division*, pop-1* expression was maintained at similar levels between daughter cells (Figure 4A-ii, B). Once the L2 symmetric division was completed, anterior daughter cells (pa) exhibited significantly higher *pop-1* mRNA compared to posterior daughters (pp) (Figure 4A-iii, B). Prior to the L2 asymmetric division, *pop-1* expression levels increased dramatically relative to the preceding symmetric division stage (Figure 4A-iv, B) and the difference between anterior-posterior cells (pa vs pp) was maintained. During anaphase-telophase of the L2 asymmetric division, *pop-1* mRNA appeared slightly enriched in posterior daughters (Figure 4A-v, B). However, following completion of the L2 asymmetric division, posterior daughter cells (pap and ppp) showed markedly lower *pop-1* mRNA levels compared to anterior daughters (paa and ppa), with 33% of cells showing no *pop-1* mRNAs (Figure 4A-vi, B), suggesting strong repression in posterior cells.

**Figure 4.**
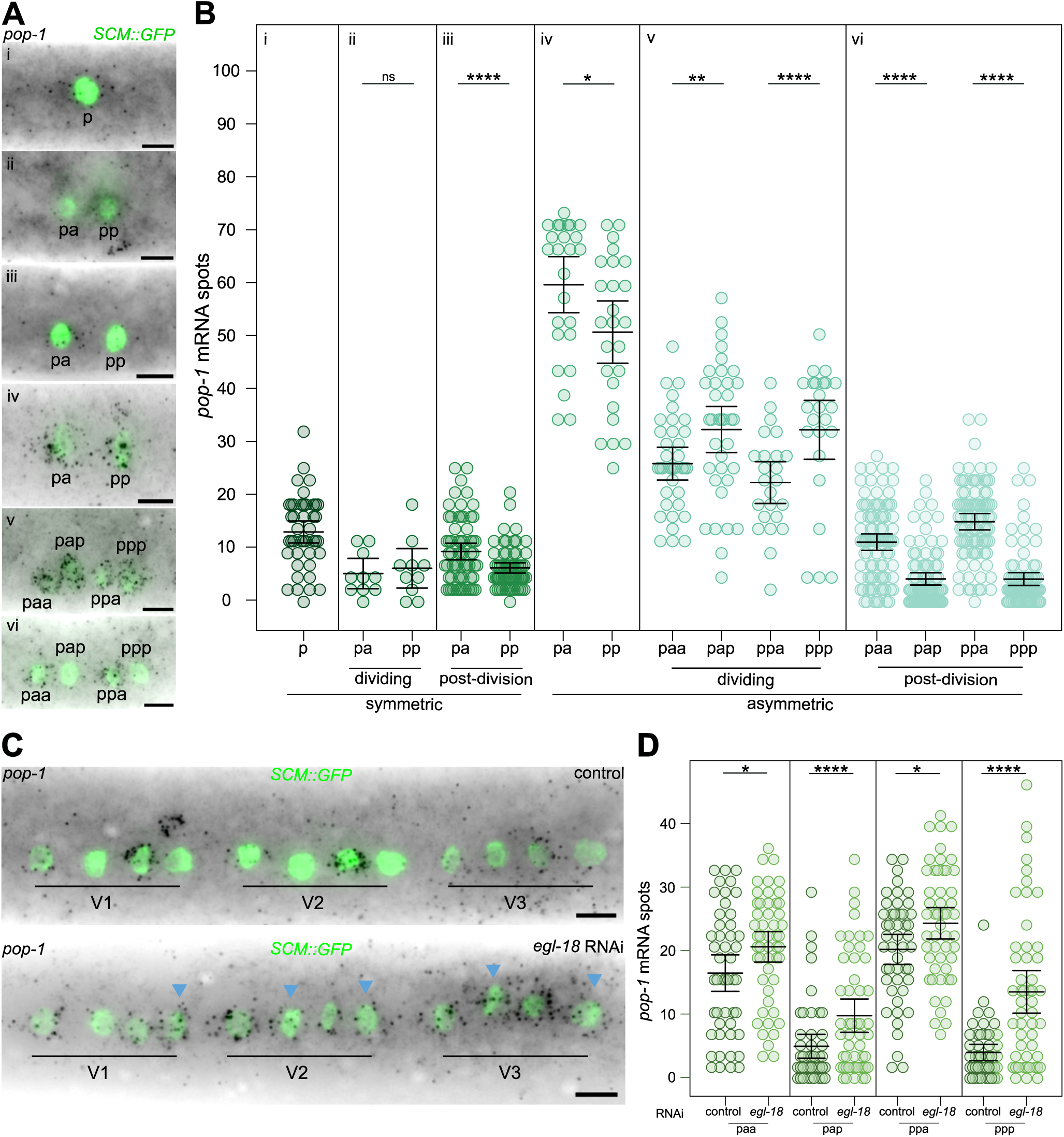
*pop-1* expression is enriched in anterior daughter cells. **(A)** Representative *pop-1* smFISH images from the late L1 stage to after the L2 asymmetric division. i- late L1, ii- during L2 symmetric division, iii- after L2 symmetric division, iv- before L2 asymmetric division, v- anaphase-telophase of L2 asymmetric division, and vi- after the L2 asymmetric division. **(B)** Quantification of *pop-1* mRNA spots in V1-V4 lineages in each of the developmental stages shown in (A) (24≤ n ≤87). Following symmetric and asymmetric divisions, anterior daughter cells (pa, paa and ppa) showed higher number of *pop-1* mRNA spots compared to corresponding posterior daughter cells (pp, ppa and ppp). **(C)** Representative *pop-1* smFISH images following the L2 asymmetric division upon *egl-18* RNAi vs control treatment. Blue arrowheads point to posterior daughter cells in *egl-18* RNAi-treated animals showing increased levels of *pop-1* transcripts compared to control. **(D)** Quantification of *pop-1* mRNA spots in V1-V4 lineages in anterior and posterior daughter cells following the L2 asymmetric division upon *egl-18* RNAi vs control treatment. Error bars in B and D show the mean ± standard deviation and **** represent *p*<0.001, ** *p*<0.01 and * p<0.05 with a two-tailed t-test. In A and C seam cells are labelled using *SCM::GFP* and scale bars are 5 μm.

We hypothesised that the differences we observed in Wnt machinery expression between anterior and posterior cells may reflect feedback from Wnt signalling activation. To test this hypothesis, we knocked down the Wnt downstream effector *egl-18* using RNAi, which leads to failures in Wnt signalling activation and decreases the terminal seam cell number (Ferrando-Marco & Barkoulas, 2025; Gorrepati et al., 2013). *egl-18* RNAi-treated animals showed a significant increase in *pop-1* expression in both anterior and posterior daughter cells with the increase in posterior cells being more striking (Figure 4C and D). These results suggest that the asymmetric *pop-1* expression pattern is indeed dependent on Wnt signalling pathway activation.

### Daughter cells resulting from symmetric and asymmetric divisions show different levels of PRY, CAM-1 and Wnt signalling activity

Next, we asked whether the unanticipated differential expression of Wnt signalling components following symmetric and asymmetric division may influence distributions at the protein level. We first chose to examine PRY-1 protein localisation using a PRY-1::mNeonGreen knock-in strain (Heppert et al., 2018), as differences in *pry-1* mRNA expression in cell daughters following symmetric and asymmetric divisions were among the most striking observed (Figure 2B). We found that prior to the L2 asymmetric division, PRY-1::mNeonGreen showed anterior membrane localisation in both pa and pp cells as expected, but pp cells displayed higher cytoplasmic PRY-1 compared to pa cells (Figure 5A-D). Following the L2 asymmetric division, daughter cells from the posterior branch (ppa and ppp) displayed significantly higher PRY-1::mNeonGreen fluorescence intensity in both the membrane and cytoplasm compared to the daughters of the anterior branch (paa and pap) (Figure 5B-D). In addition, posterior daughter cells (pap and ppp) showed significantly higher levels of cytoplasmatic PRY-1 compared to their respective posterior daughter cell (paa and ppa), which is consistent with the smFISH data.

**Figure 5.**
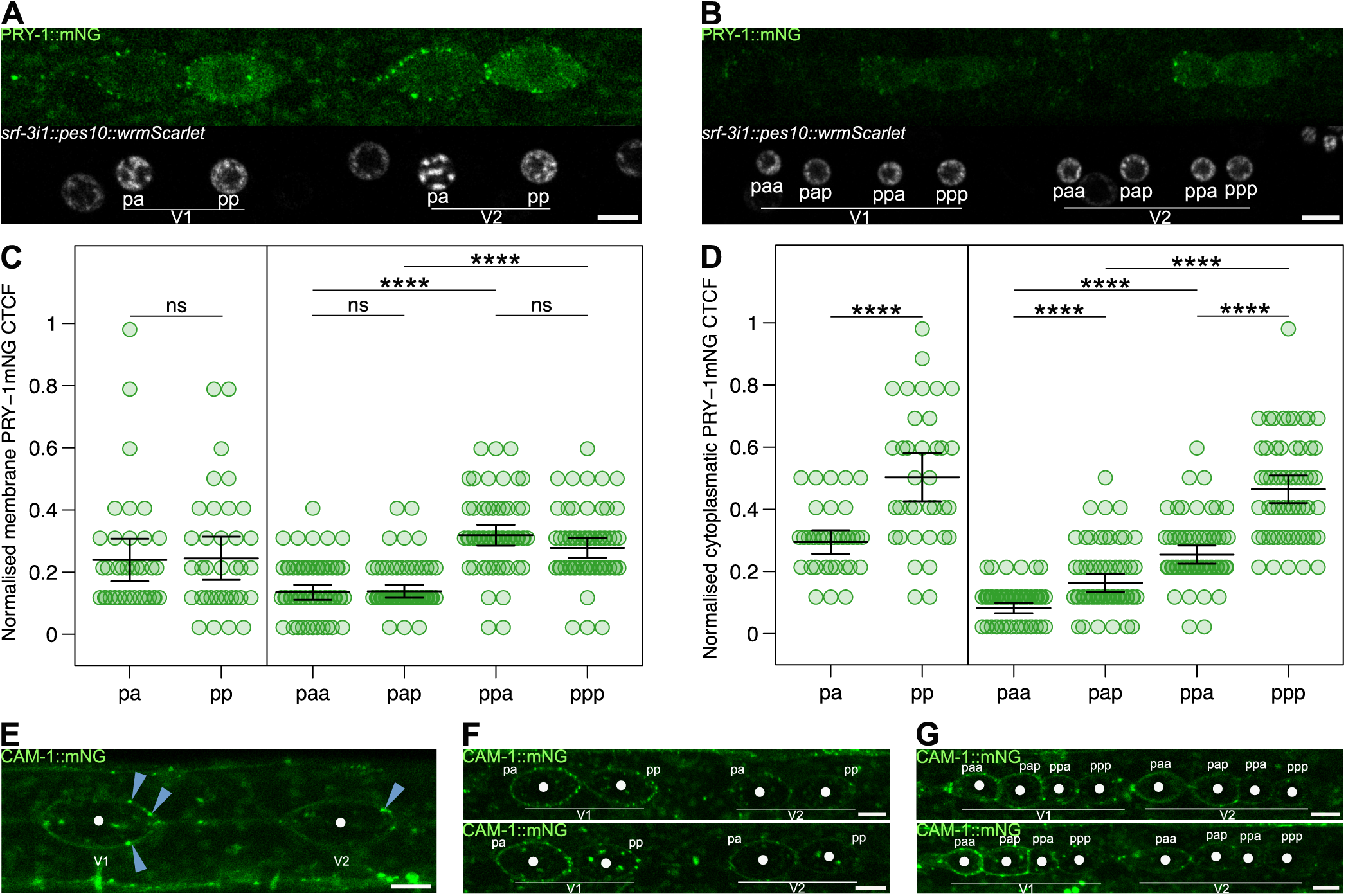
Analysis of PRY-1 and CAM-1 expression following symmetric and asymmetric divisions shows asymmetric protein localisation between daughter cells. **(A-B)** Representative PRY-1::mNeonGreen images before (A) and after (B) the L2 asymmetric division. Seam cell nuclei is labelled with *srf-3i::pes-10::wrmScarlet*. **(C-D)** Quantification of PRY-1::mNeonGreen fluorescence intensity following the L2 symmetric (pa, pp) and asymmetric (paa, pap, ppa, ppp) divisions in the membrane (C) and in the cytoplasm (D). Error bars show the mean ± standard deviation and **** represent *p*<0.001 with a two-tailed t-test, 36 ≤ n ≤ 60 cells per condition **(E-G)** Representative CAM-1::mNeonGreen images at late L1 (E) and following the L2 symmetric (F) and asymmetric (G) divisions. White dots mark the approximate centre of each cell to aid identification. Blue arrows indicate posterior membrane-localised foci. Scale bars are 5 μm in A, B, E, F and G.

We then focused on the expression of the Ror receptor CAM-1, for which we previously reported that *cam-1* mRNA is enriched in anterior daughter cells following both symmetric and asymmetric divisions (Ferrando-Marco et al., 2026). We used a CAM-1::mNeonGreen knock-in strain (Heppert et al., 2018) to study protein localisation. At late L1, mother cells showed posterior membrane localisation as previously reported for other Wnt receptors (Mizumoto & Sawa, 2007b, 2007a). We focused on V1 cells because, across all animals analysed, CAM-1 signal was consistently stronger in the V1 lineage compared to V2-V4, in agreement with what we previously reported for *cam-1* mRNA (Ferrando-Marco et al., 2026). Following symmetric division, 57% animals analysed showed a clear enrichment of CAM-1 at the posterior cortex of anterior daughter cells (pa) compared to posterior daughter cells in the V1 lineage (Figure 5F). Following asymmetric division, 80% animals analysed displayed higher membrane-localised CAM-1 in the anterior branch (paa and pap) relative to daughter cells from the posterior branch (ppa and ppp) (Figure 5G), which is consistent with the smFISH data.

The differential expression of Wnt components in the anterior and posterior branches within each lineage raised the possibility that this might stem from or result in differences in the levels of activation of the Wnt signalling pathway in daughter cells. To test this hypothesis, we utilised the POPHHOP (POP-1 and HMG-helper optimal promoter) reporter, which serves as a readout of Wnt signalling activation but has not been imaged before in the context of dividing seam cells (Bhambhani et al., 2014; Katsanos et al., 2017). We performed time-lapse imaging using microfluidic chambers to track POPHHOP reporter expression dynamics (Berger et al., 2021). Time-lapse imaging analysis spanning from late L1 (t=1) through late L2 (t=36) revealed dynamic POPHHOP expression patterns (Figure 6A). At late L1, POPHHOP expression was detectable in p cells. Following the L2 symmetric division, pp cells showed higher POPHHOP activity compared to pa cells in 58% of pairs visualised. Consistent with this, cells derived from the posterior branch (ppa and ppp) displayed higher POPHHOP expression frequencies than those from the anterior branch (paa and pap) in 68% of cases (Figure 6B). Given the high stability of the GFP reporter and the short time interval between divisions, GFP signal detected in later daughter cells likely reflects inheritance from the pp cell where activation is higher than pa, perhaps above a threshold allowing GFP detection. Taken together, these results suggest that differential expression of Wnt pathway components may be associated with differences in Wnt signalling pathway activity between pa and pp daughters following symmetric division, mediated by feedback mechanisms.

**Figure 6.**
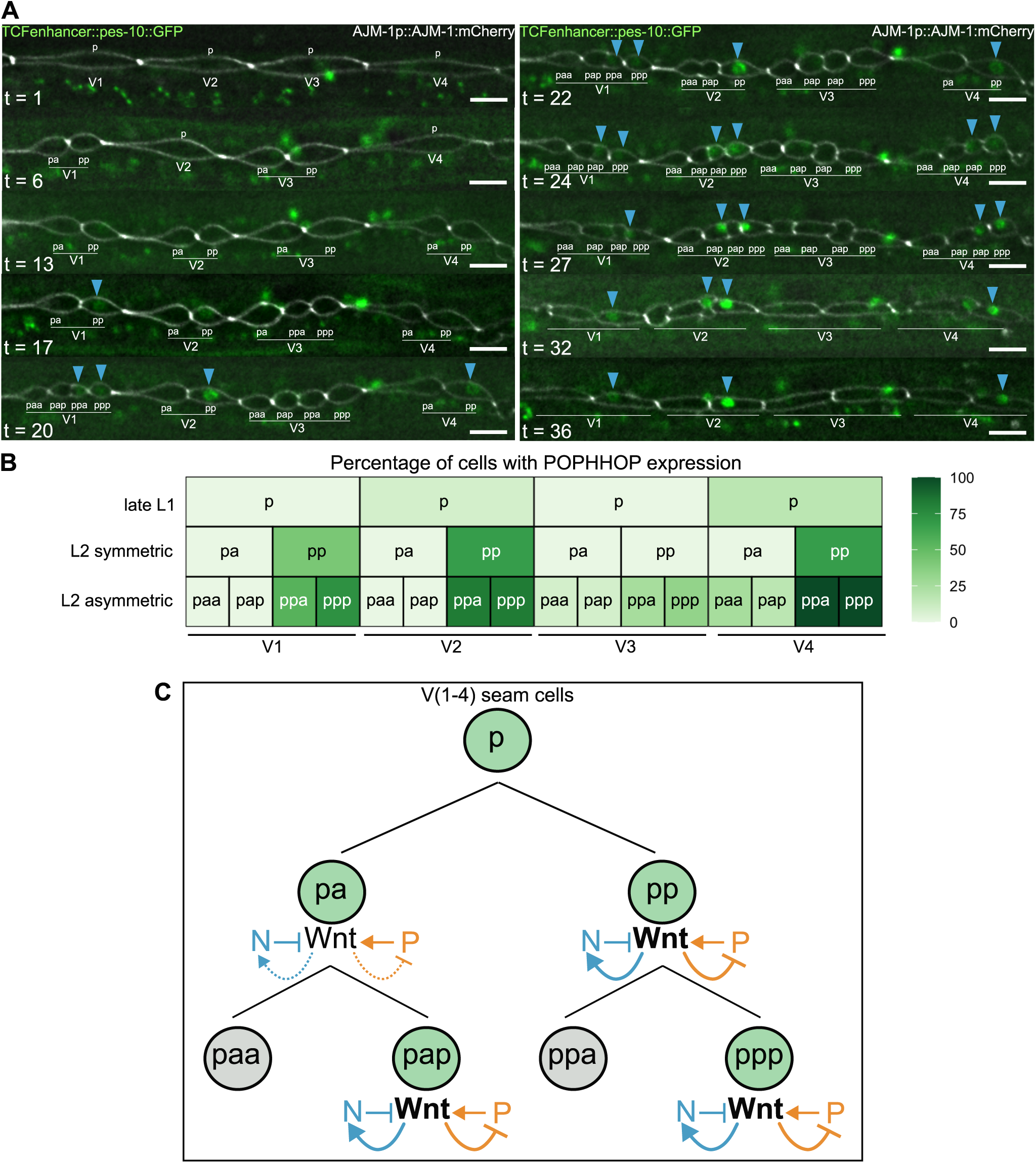
Daughter cells resulting from symmetric division show differential Wnt activation capacity. **(A)** Representative images of POPHHOP (POP-1 and HMG-helper optimal promoter) reporter signal form late L1 (t=1) to late L2 stage (t=36). Time between frames (t) is 15 minutes. The membrane of the seam cells is labelled using the apical junction marker *ajm-1p::ajm-1::mCherry.* Blue arrows point to cells with GFP expression. Scale bars are 10 μm. **(B)** Heatmap showing percentage of cells with GFP expression in V1-V4 lineages at late L1 (p), and following the L2 symmetric (pa and pp) and asymmetric divisions (paa, pap, ppa, ppp). Cells in the posterior branch of the lineage (pp, ppa and ppp) show stronger POPHHOP expression compared to cells in the anterior branch (pa, paa and pap). n = 12 animals. **(C)** Schematic representation of proposed feedback regulation within the Wnt signalling pathway across symmetric and asymmetric seam cell divisions. Wnt signalling activation is hypothesised to regulate the transcription of both negative (N) and positive (P) pathway components. Upregulation of negative regulators and repression of positive regulators may act as feedback mechanisms to attenuate pathway activity following fate specification, contributing to robust asymmetric divisions. Following the symmetric division, posterior daughter cells are proposed to exhibit stronger Wnt-dependent feedback compared to anterior daughters, consistent with higher Wnt signalling activity in these cells.

## Discussion

Comprehensive analysis of mRNA distribution patterns of Wnt pathway components following L2 symmetric and asymmetric division revealed enrichment for the negative regulators *pry-1* and *apr-1* in posterior daughter cells and enrichment for the positive regulators *sys-1*, *wrm-1*, and *lit-1*, along with the transcription factor *pop-1* in anterior daughter cells. These expression patterns are intriguing because they appear counter-intuitive given what is known about Wnt pathway activation in the context of the Wnt asymmetry model. Furthermore, the mRNA distributions we report during the L2 divisions were not anticipated based on the previously reported protein distributions based on translational reporters. For example, PRY-1 and APR-1 have been shown to localise to the anterior cortex at the L1 stage during division (Mizumoto & Sawa, 2007a, 2007b), whilst WRM-1, SYS-1 and LIT-1 show posterior nuclear enrichment in daughter cells (Mizumoto & Sawa, 2007b; Takeshita & Sawa, 2005). In light of the smFISH results, we revisited the expression of a PRY-1 reporter during the L2 divisions and found that the protein accumulation pattern correlated well with transcript quantification. Moreover, the *pop-1* mRNA pattern is consistent with the known protein localisation. POP-1 is a transcription factor whose activity is primarily controlled by nuclear export upon phosphorylation (Lo et al., 2004; Meneghini et al., 1999; Rocheleau et al., 1999; Yang et al., 2011). Since this phosphorylation is favoured in posterior daughter cells, this leads to anterior nuclear enrichment (Calvo et al., 2001; Phillips et al., 2007; Shetty et al., 2005), which is consistent with *pop-1* mRNA being enriched in anterior daughters following asymmetric division.

We reasoned that the transcriptional patterns reported here might be a consequence of feedback within the Wnt signalling pathway. For example, Axin is a known target of Wnt signalling in vertebrates and *Drosophila* (Jho et al., 2002; Lustig et al., 2001), and in *C. elegans pry-1* is upregulated upon Wnt signalling activation (Gorrepati & Eisenmann, 2015; Jackson et al., 2014). We formally tested this idea for *pop-1* regulation by showing that its asymmetric expression is disrupted when *egl-18*, a target of the Wnt signalling (Gorrepati et al., 2013; Gorrepati & Eisenmann, 2015), is downregulated. This suggests that *pop-1* repression in posterior daughter cells might be reinforced by Wnt signalling, and this feedback might contribute to robust commitment to seam cell fate following asymmetric division. Transcriptional regulation of TCF/LEF by the Wnt signalling pathway has been described before. For instance, in colon cancer cells LEF1 transcription is activated by β-catenin/TCF complexes (Cadigan & Waterman, 2012; Hovanes et al., 2001; T. W. H. Li et al., 2006). Similarly, β-catenin and TCF4 activate TCF1 expression in intestinal epithelial cells (Roose et al., 1998). However, in human embryonic kidney cells, Wnt-dependent regulation of LEF1 expression has been shown to occur independently of TCF/LEF-β-catenin complexes, possibly involving other Wnt-induced transcription factors (Filali et al., 2002). It is possible that the reported transcriptional regulation of Wnt components may also reflect direct input from cell fate determination programmes and is independent to some extent of Wnt signalling.

Feedback regulation might extend to other members of the Wnt signalling pathway. For example, we have previously described differential expression of Wnt receptors and *Dishevelled* between the daughter cells following symmetric and asymmetric divisions, with *lin-17* being enriched in posterior daughters and *cam-1*, *mom-5* and *dsh-2* being enriched in anterior daughters, potentially as a result of feedback regulation (Ferrando-Marco et al., 2026). Both positive and negative regulation of Wnt receptors has been reported in *C. elegans* in the case of Q neuroblasts, where Wnt signalling promotes *lin-17* expression while it represses *mom-5* expression (Ji et al., 2013). Regulation of Frizzled receptor expression by Wnt signalling has also been described in other systems, including *Drosophila* and human embryonic stem cells, suggesting that receptor-level feedback is a common feature of Wnt pathway regulation (Cardigan et al., 1998; Sato et al., 1999; Willert et al., 2002).

What is the need for such extensive transcriptional feedback? The feedback in the Wnt pathway may help ensure robust asymmetric divisions, in line with the pervasive role of feedback regulation in biological systems (Félix & Barkoulas, 2015; Masel & Siegal, 2009; Whitacre, 2012). For example, repression of negative regulator such as PRY-1/Axin in anterior cells may safeguard cell fate allocation during asymmetric division while enrichment in posterior cells may help attenuate pathway activity after initial fate specification, preventing excessive signalling. Similarly, transcriptional repression of positive regulators such as β-catenins in posterior daughter cells following asymmetric division may also contribute to priming the system for subsequent divisions. Successive posterior enrichment and accumulation of β-catenin in asymmetric divisions has been discussed as a potential problem in the context of Wnt signalling activity repression in anterior daughter cells (Lam & Phillips, 2017). Transcriptional repression in posterior daughters might be important to limit SYS-1 and WRM-1 levels ensuring that when cells subsequently divide asymmetrically, their anterior daughters inherit lower levels of β-catenin protein, facilitating efficient degradation by the destruction complex. This transcriptional mechanism would complement post-translational regulatory mechanisms, including centrosome localisation and cell cycle-coupled degradation, to ensure robust asymmetric cell fate decisions (Vora & Phillips, 2015).

We report distinct capacities for Wnt signalling activation between the daughters of symmetric divisions, indicating that pa and pp cells are not identical despite adopting the same fate (Figure 6C). This finding suggests that the Wnt-dependent transcriptional feedback mechanisms that shape asymmetric division outcomes may also operate at the symmetric division stage. The differences in Wnt pathway activation following symmetric division are likely rooted in the polarity established in the mother cell, leading to asymmetric inheritance of Wnt activation potential in the daughters. Notably, high-Wnt cells are enriched for negative regulators, whereas low-Wnt cells are enriched for positive regulators. This pattern indicates that the observed mRNA distributions more likely reflect differential Wnt pathway activation in the daughter cells and associated feedback mechanisms, rather than being the primary cause of these differences. Wnt pathway components such as POP-1, WRM-1, LIT-1 have been previously shown to be asymmetrically localised even during symmetric divisions (Harandi & Ambros, 2015; Hughes et al., 2013; van der Horst et al., 2019; Wildwater et al., 2011; Ferrando-Marco & Barkoulas, 2025). In agreement with these studies, we find that in nearly all cases, the expression of Wnt machinery differs between daughter cells following symmetric division at the transcript and protein level. The molecular heterogeneity between the anterior and posterior lineage branches may have functional consequences, particularly in response to genetic or environmental perturbations. Indeed, the anterior branch has been reported to exhibit increased sensitivity to temperature shifts and mutations (Hintze et al., 2020; van der Horst et al., 2019). Regardless of the precise functional implications, our findings suggest that seam cell lineages may be less homogeneous than previously assumed. Even divisions that appear symmetric give rise to daughter cells with clear molecular asymmetries, which is important to be considered in the analysis and interpretation of developmental phenotypes.

## Materials and Methods

### *C. elegans* maintenance and genetics

All *C. elegans* strains in this study were maintained at 20°C on Nematode Growth Medium (NGM) plates seeded with *E. coli* OP50 (Brenner, 1974). A list of strains used in this study is provided in Table S1.

### RNAi feeding

RNAi by feeding was performed using *E. coli* HT115 expressing double-stranded RNA targeting *egl-18*, as previously described (Kamath et al., 2003; Kamath & Ahringer, 2003). Bacterial cultures were grown overnight, and 300 μL were seeded onto NGM plates containing 25 μg/mL ampicillin, 12.5 μg/mL tetracycline, and 1 mM IPTG. RNAi treatment was performed during postembryonic development by seeding eggs of the indicated strains directly onto *egl-18* RNAi plates. The *egl-18* RNAi clone used in this study was obtained from the Ahringer RNAi Library (Kamath & Ahringer, 2003) (Source Bioscience).

### Single molecule fluorescent in situ hybridisation

Animals were synchronised by bleaching and were fixed at 25h post-bleaching 4% formaldehyde for 45 minutes and permeabilised with 70% ethanol. Samples were then incubated with a pool of 24-48 oligos labelled with Quasar 670 (Biosearch Technologies, Novato, CA) for 16 hours. Imaging was performed using the x100 objective of an epifluorescence Ti-eclipse microscope (Nikon) and DU-934 CCD-17291 camera (Andor Technology) operated via the Nikon NIS Elements Software. 23 z-stack slices of 0.8 μm were acquired for each animal. Spot quantification was performed using MATLAB (MathWorks) as previously described (Raj et al., 2008). Z-slices containing spots where max-projected to give a single image, inverted and merged to the GFP channel using ImageJ (NIH). A complete list of the smFISH probes used in this study is provided in Table S1.

### Confocal microscopy

Protein reporter images were acquired using a Leica SP8 Inverted confocal microscope with a x63/1.4 oil DIC objective controlled by LAS X software. Animals were mounted in 5% agarose pads containing 100 μM sodium azide (NaN_3_) and secured with a coverslip. To visualise L2 symmetric and asymmetric divisions animals were observed at 26 hours post-bleaching. Z-stack slices of 0.4 μm were acquired and 7 slices were SUM projected for quantification using Fiji/ImageJ. For each seam-cell daughter, membrane-associated PRY-1::mNeonGreen signal was quantified using a segmented line selection manually traced along the cell cortex. The line was drawn to follow the membrane and set to a width sufficient to fully encompass the visible membrane-associated PRY-1 signal, including punctate cortical accumulations, while minimising cytoplasmic contribution. Cytoplasmic PRY-1::mNeonGreen signal was quantified using a manually defined area selection drawn within the cell body. Cytoplasmic regions were selected based on cell morphology without explicit nuclear exclusion. For both membrane and cytoplasmic measurements, fluorescence intensity was quantified as corrected total cell fluorescence (CTCF), calculated as:

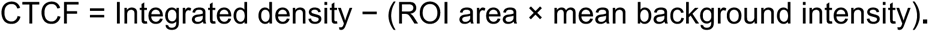

Background fluorescence was measured for each image using three regions of interest placed in signal-free areas outside the animal, and the average background value was used for correction. All images were acquired and analysed using identical settings.

The data was normalised (rescaled) to [0,1] using the formula:

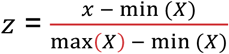

where X is the data, x is a datapoint and z is the corresponding rescaled datapoint. Picture editing was performed using straightening tool and stitching macro from FIJI (Preibisch et al., 2009).

### Microfluidic imaging

Microfluidic imaging was performed according to (Berger et al., 2021, 2025). Briefly, NA22 bacteria were grown overnight in 40 mL LB with shaking at 37 °C to an OD600 of approx. 1.9, washed three times with S-basal, and finally resuspended in approx. 1 mL S-basal. Finally, 1 mL of bacterial suspension was mixed with 0.65 mL Optiprep density gradient medium and 0.33 mL S-basal supplemented with 1% Pluronic F127, and finally filtered using a 10 µm strainer.

Animals were synchronized by bleaching, and embryos were allowed to hatch overnight in M9, arresting at the L1 larval stage. For imaging at late L1, animals were plated on NGM plates, grown for 16 hours, and collected using S-basal.

Animals were then loaded into an L1-L4 microfluidic device and imaged at 15-minute intervals for 24 hours using an upright microscope (DMRA2, Leica), equipped with a sCMOS camera (Prime BSI, Photometrics), a LED combiner (Spectra, Lumencor) and a piezo objective drive (MIPOS 100, Piezosystems Jena). Z-stacks of 0.5 μm were acquired using a x40 oil objective. Image acquisition was controlled using custom-built MATLAB scripts (MATLAB 2023b, MathWorks) and custom-built microcontrollers (Arduino Mega 2560) for coordination of fluorescence and brightfield LEDs, piezo and camera. All images were acquired at 20±0.5°C, with temperature control achieved either via a microscope cage incubator (H201-T-UNIT-BL-CRYO and H201-ENCLOSURE-CRYO, Okolab).

Images were deconvolved using using the YacuDecu implementation of CUDA-based Richardson Lucy deconvolution in MATLAB (www.github.com/bobpepin/YacuDecu) with PSFs generated using microPSF (J. Li et al., 2017) and images were edited for presentation using Fiji/ImageJ.

### Quantification and statistical analysis

Graphic representation and statistical analysis were performed using R. Error bars used in all graphs represent the standard deviation. An unpaired t-test was used to evaluate significance as indicated in the figure legends. Results were considered statistically significant when p < 0.05. Asterisks in figures indicate corresponding statistical significance as it follows: * p < 0.05; ** p < 0.01; *** p < 0.005; **** p < 0.001.

### Contact for Reagent and Resource Sharing

Further information and requests for resources and reagents should be directed to and will be fulfilled by the Lead Contact, Michalis Barkoulas.

## Acknowledgements

This work was supported by the Wellcome Trust [219448/Z/19/Z]. We thank Alex Hanjal for supporting this work in his lab. Some strains were provided by the CGC, which is funded by NIH Office of Research Infrastructure Programs (P40 OD010440). We also thank the facility for Imaging by Light Microscopy (FILM) at Imperial College London, which is part-supported by funding from the Wellcome Trust [104931/Z/14/Z] and the BBSRC [BB/T017929/1].

## Author Contributions

MFM carried out experiments and data analysis. SB carried out microfluidics imaging. MB supervised the work. MFM and MB wrote the manuscript.

## References

Baldwin, A. T., & Phillips, B. T. (2014). The tumor suppressor APC differentially regulates multiple β-catenins through the function of axin and CKIα during *C. elegans* asymmetric stem cell divisions. Journal of Cell Science, 127(12), 2771–2781. 10.1242/jcs.146514

Banerjee, D., Chen, X., Lin, S. Y., & Slack, F. J. (2010). *kin-19*/casein kinase Iα has dual functions in regulating asymmetric division and terminal differentiation in *C. elegans* epidermal stem cells. Cell Cycle, 9(23), 4748–4765. 10.4161/cc.9.23.14092

Bekas, K. N., & Philips, B. T. (2022). *unc-37*/Groucho and *lsy-22*/AES repress Wnt target genes in *C. elegans* asymmetric cell divisions. BioRxiv. 10.1101/2022.01.10.475695

Berger, S., Spiri, S., deMello, A., & Hajnal, A. (2021). Microfluidic-based imaging of complete *Caenorhabditis elegans* larval development. Development (Cambridge), 148(18). 10.1242/DEV.199674

Berger, S., Spiri, S., Demello, A., & Hajnal, A. (2025). High-Resolution *C. elegans* Imaging Across All Larval Stages. Journal of Visualized Experiments, 2025-May(219). 10.3791/68172

Bhambhani, C., Ravindranath, A. J., Mentink, R. A., Chang, M. V., Betist, M. C., Yang, Y. X., Koushika, S. P., Korswagen, H. C., & Cadigan, K. M. (2014). Distinct DNA Binding Sites Contribute to the TCF Transcriptional Switch in *C. elegans* and *Drosophila*. PLoS Genetics, 10(2). 10.1371/journal.pgen.1004133

Bielen, H., & Houart, C. (2014). The Wnt cries many: Wnt regulation of neurogenesis through tissue patterning, proliferation, and asymmetric cell division. Developmental Neurobiology, 74(8), 772–780. 10.1002/dneu.22168

Boukhibar, L. M., & Barkoulas, M. (2016). The developmental genetics of biological robustness. Annals of Botany, 117(5), 699–707. 10.1093/aob/mcv128

Brenner, S. (1974). The genetics of *Caenorhabditis elegans*. Genetics, 77(1), 71–94. 10.1093/genetics/77.1.71

Cadigan, K. M., & Waterman, M. L. (2012). TCF/LEFs and Wnt signaling in the nucleus. Cold Spring Harbor Perspectives in Biology, 4(11). 10.1101/cshperspect.a007906

Calvo, D., Victor, M., Gay, F., Sui, G., Po-Shan Luke, M., Dufourcq, P., Wen, G., Maduro, M., Rothman, J., & Shi, Y. (2001). A POP-1 repressor complex restricts inappropriate cell type-specific gene transcription during *Caenorhabditis elegans* embryogenesis. EMBO Journal, 20(24), 7197–7208. 10.1093/emboj/20.24.7197

Cardigan, K. M., Fish, M. P., Rulifson, E. J., & Nusse, R. (1998). Wingless Repression of *Drosophila frizzled 2* Expression Shapes the Wingless Morphogen Gradient in the Wing. Cell, 93, 767–777.

Chisholm, A. D., & Hsiao, T. I. (2012). The *Caenorhabditis elegans* epidermis as a model skin. I: Development, patterning, and growth. Wiley Interdisciplinary Reviews: Developmental Biology, 1(6), 861–878. 10.1002/wdev.79

Félix, M. A., & Barkoulas, M. (2015). Pervasive robustness in biological systems. Nature Reviews Genetics, 16(8), 483–496. 10.1038/nrg3949

Ferrando-Marco, M., & Barkoulas, M. (2025). EFL-3/E2F7 modulates Wnt signalling by repressing the Nemo-like kinase LIT-1 during asymmetric epidermal cell division in *Caenorhabditis elega*ns. Development (Cambridge), 152(5). 10.1242/dev.204546

Ferrando-Marco, M., Garcia del Valle, B., Hintze, M., Narunsky, L., Lin, S., Huang, J., Edwards, S., & Barkoulas, M. (2026). HDAC1/2-mediated repression of Wnt receptor expression orients asymmetric division polarity in *C. elegans*. BioRxiv. 10.64898/2026.02.23.707389

Filali, M., Cheng, N., Abbott, D., Leontiev, V., & Engelhardt, J. F. (2002). Wnt-3A/β-catenin signaling induces transcription from the LEF-1 promoter. Journal of Biological Chemistry, 277(36), 33398–33410. 10.1074/jbc.M107977200

Gleason, J. E., & Eisenmann, D. M. (2010). Wnt signaling controls the stem cell-like asymmetric division of the epithelial seam cells during *C. elegans* larval development. Developmental Biology, 348(1), 58–66. 10.1016/j.ydbio.2010.09.005

Goldstein, B., Takeshita, H., Mizumoto, K., & Sawa, H. (2006). Wnt signals can function as positional cues in establishing cell polarity. Developmental Cell, 10(3), 391–396. 10.1016/j.devcel.2005.12.016

Gorrepati, L., & Eisenmann, D. M. (2015). The *C. elegans* embryonic fate specification factor EGL-18 (GATA) is reutilized downstream of Wnt signaling to maintain a population of larval progenitor cells. Worm, 4(1), e996419. 10.1080/23723556.2014.996419

Gorrepati, L., Thompson, K. W., & Eisenmann, D. M. (2013). *C. elegans* GATA factors EGL-18 and ELT-6 function downstream of Wnt signaling to maintain the progenitor fate during larval asymmetric divisions of the seam cells. Development (Cambridge), 140(10), 2093–2102. 10.1242/dev.091124

Green, J. L., Inoue, T., & Sternberg, P. W. (2008). Opposing Wnt Pathways Orient Cell Polarity during Organogenesis. Cell, 134(4), 646–656. 10.1016/j.cell.2008.06.026

Habib, S. J., & Acebrón, S. P. (2022). Wnt signalling in cell division: from mechanisms to tissue engineering. Trends in Cell Biology, 32(12), 1035–1048. 10.1016/j.tcb.2022.05.006

Harandi, O. F., & Ambros, V. R. (2015). Control of stem cell self-renewal and differentiation by the heterochronic genes and the cellular asymmetry machinery in *Caenorhabditis elegans*. PNAS, 112(3), E287–E296. 10.1073/pnas.1422852112

Heppert, J. K., Pani, A. M., Roberts, A. M., Dickinson, D. J., & Goldstein, B. (2018). A CRISPR tagging-based screen reveals localized players in Wnt-directed asymmetric cell division. Genetics, 208(3), 1147–1164. 10.1534/genetics.117.300487

Hintze, M., Koneru, S. L., Gilbert, S. P. R., Katsanos, D., Lambert, J., & Barkoulas, M. (2020). A cell fate switch in the *Caenorhabditis elegans* seam cell lineage occurs through modulation of the wnt asymmetry pathway in response to temperature increase. Genetics, 214(4), 927–939. 10.1534/GENETICS.119.302896

Hovanes, K., Li, T. W. H., Munguia, J. E., Truong, T., Milovanovic, T., Marsh, J. L., Holcombe, R. F., & Waterman, M. L. (2001). β-catenin-sensitive isoforms of lymphoid enhancer factor-1 are selectively expressed in colon cancer. Nature Genetics, 28, 53–57. 10.1038/ng0501-53

Huang, S., Shetty, P., Robertson, S. M., & Lin, R. (2007). Binary cell fate specification during *C. elegans* embryogenesis driven by reiterated reciprocal asymmetry of TCF POP-1 and its coactivator β-catenin SYS-1. Development, 134(14), 2685–2695. 10.1242/dev.008268

Hughes, S., Brabin, C., Appleford, P. J., & Woollard, A. (2013). CEH-20/Pbx and UNC-62/Meis function upstream of *rnt-1*/Runx to regulate asymmetric divisions of the *C. elegans* stem-like seam cells. Biology Open, 2(7), 718–727. 10.1242/bio.20134549

Jackson, B. M., Abete-Luzi, P., Krause, M. W., & Eisenmann, D. M. (2014). Use of an activated beta-catenin to identify wnt pathway target genes in Caenorhabditis elegans, including a subset of collagen genes expressed in late larval development. G3: Genes, Genomes, Genetics, 4(4), 733–747. 10.1534/g3.113.009522

Jho, E., Zhang, T., Domon, C., Joo, C.-K., Freund, J.-N., & Costantini, F. (2002). Wnt/β-Catenin/Tcf Signaling Induces the Transcription of Axin2, a Negative Regulator of the Signaling Pathway. Molecular and Cellular Biology, 22(4), 1172–1183. 10.1128/mcb.22.4.1172-1183.2002

Ji, N., Middelkoop, T. C., Mentink, R. A., Betist, M. C., Tonegawa, S., Mooijman, D., Korswagen, H. C., & Van Oudenaarden, A. (2013). Feedback control of gene expression variability in the *Caenorhabditis elegans* Wnt pathway. Cell, 155(4), 869. 10.1016/j.cell.2013.09.060

Joshi, P. M., Riddle, M. R., Djabrayan, N. J. V., & Rothman, J. H. (2010). *Caenorhabditis elegans* as a model for stem cell biology. Developmental Dynamics, 239(5), 1539–1554. 10.1002/DVDY.22296

Kamath, R. S., & Ahringer, J. (2003). Genome-wide RNAi screening in *Caenorhabditis elegans*. Methods, 30(4), 313–321. 10.1016/S1046-2023(03)00050-1

Kamath, R. S., Fraser, A. G., Dong, Y., Poulin, G., Durbin, R., Gotta, M., Kanapin, A., Le Bot, N., Moreno, S., Sohrmann, M., Welchman, D. P., Zipperien, P., & Ahringer, J. (2003). Systematic functional analysis of the *Caenorhabditis elegans* genome using RNAi. Nature, 421(6920), 231–237. 10.1038/nature01278

Katsanos, D., Koneru, S. L., Mestek Boukhibar, L., Gritti, N., Ghose, R., Appleford, P. J., Doitsidou, M., Woollard, A., van Zon, J. S., Poole, R. J., & Barkoulas, M. (2017). Stochastic loss and gain of symmetric divisions in the *C. elegans* epidermis perturbs robustness of stem cell number. PLoS Biology, 15(11), 1–31. 10.1371/journal.pbio.2002429

Knoblich, J. A. (2008). Mechanisms of Asymmetric Stem Cell Division. Cell, 132(4), 583–597. 10.1016/j.cell.2008.02.007

Koneru, S. L., Hintze, M., Katsanos, D., & Barkoulas, M. (2021). Cryptic genetic variation in a heat shock protein modifies the outcome of a mutation affecting epidermal stem cell development in *C. elegans*. Nature Communications, 12(1), 1–12. 10.1038/s41467-021-23567-1

Lam, A. K., & Phillips, B. T. (2017). Wnt signaling polarizes *C. elegans* asymmetric cell divisions during development. Results and Problems in Cell Differentiation, 61, 83–114. 10.1007/978-3-319-53150-2_4

Li, J., Xue, F., & Blu, T. (2017). Fast and accurate three-dimensional point spread function computation for fluorescence microscopy. Journal of the Optical Society of America A, 34(6), 1029. 10.1364/josaa.34.001029

Li, T. W. H., Ting, J.-H. T., Yokoyama, N. N., Bernstein, A., van de Wetering, M., & Waterman, M. L. (2006). Wnt Activation and Alternative Promoter Repression of LEF1 in Colon Cancer. Molecular and Cellular Biology, 26(14), 5284–5299. 10.1128/mcb.00105-06

Lo, M.-C., Gay, F., Odom, R., Shi, Y., & Lin, R. (2004). Phosphorylation by the-Catenin/MAPK Complex Promotes 14-3-3-Mediated Nuclear Export of TCF/POP-1 in Signal-Responsive Cells in *C. elegans*. Cell, 117, 95–106. 10.1016/s0092-8674(04)00203-x

Lustig, B., Jerchow, B., Sachs, M., Weiler, S., Pietsch, T., Rarsten, U., Van De Wetering, M., Clevers, H., Schlag, P. M., Birchmeier, W., & Behrens, J. (2001). Negative feedback loop of Wnt signaling through upregulation of conductin/axin2 in colorectal and liver tumors. Langenbeck’s Archives of Surgery, 386(6), 466. 10.1128/mcb.22.4.1184-1193.2002

Masel, J., & Siegal, M. L. (2009). Robustness: mechanisms and consequences. Trends in Genetics, 25(9), 395–403. 10.1016/j.tig.2009.07.005

Meneghini, M. D., Ishitani, T., Carter, C. J., Hisamoto, N., Ninomiya-Tsuji, J., Thorpe, C. J., Hamill, D. R., Matsumoto, K., & Bowerman, B. (1999). MAP kinase and Wnt pathways converge to downregulate an HMG-domain repressor in *Caenorhabditis elegans*. Nature, 399, 793–797.

Mizumoto, K., & Sawa, H. (2007a). Cortical β-Catenin and APC Regulate Asymmetric Nuclear β-Catenin Localization during Asymmetric Cell Division in *C. elegans*. Developmental Cell, 12(2), 287–299. 10.1016/j.devcel.2007.01.004

Mizumoto, K., & Sawa, H. (2007b). Two βs or not two βs: regulation of asymmetric division by β-catenin. Trends in Cell Biology, 17(10), 465–473. 10.1016/J.TCB.2007.08.004

Morrison, S. J., & Kimble, J. (2006). Asymmetric and symmetric stem-cell divisions in development and cancer. Nature, 441(7097), 1068–1074. 10.1038/nature04956

Phillips, B. T., III, A. R. K., King, R., Hardin, J., & Kimble, J. (2007). Reciprocal asymmetry of SYS-1/β-catenin and POP-1/TCF controls asymmetric divisions in *Caenorhabditis elegans*. PNAS, 104(9), 3231–3236. 10.1073/pnas.0611507104

Phillips, B. T., & Kimble, J. (2009). A New Look at TCF and β-Catenin through the Lens of a Divergent *C. elegans* Wnt Pathway. Developmental Cell, 17(1), 27–34. 10.1016/j.devcel.2009.07.002

Preibisch, S., Saalfeld, S., & Tomancak, P. (2009). Globally optimal stitching of tiled 3D microscopic image acquisitions. Bioinformatics, 25(11), 1463–1465. 10.1093/bioinformatics/btp184

Raj, A., van den Bogaard, P., Rifkin, S. A., van Oudenaarden, A., & Tyagi, S. (2008). Imaging individual mRNA molecules using multiple singly labelled probes. Nature Methods, 5(10), 877–879. 10.1038/nmeth.1253

Rocheleau, C. E., Yasuda, J., Shin, T. H., Lin, R., Sawa, H., Okano, H., Priess, J. R., Davis, R. J., & Mello, C. C. (1999). WRM-1 Activates the LIT-1 Protein Kinase to Transduce Anterior/Posterior Polarity Signals in *C. elegans*. Cell, 97, 717–726. 10.1016/s0092-8674(00)80784-9

Roose, J., Huls, G., van Beest, M., Moerer, P., van der Horn, K., Goldschmeding, R., Logtenberg, T., & Clevers, H. (1998). Synergy Between Tumor Suppressor APC and the-Catenin-Tcf4 Target Tcf1. Science, 281, 1923–1926. 10.1126/science.285.5435.1923

Sato, A., Kojima, T., Ui-Tei, K., Miyata, Y., & Saigo, K. (1999). Dfrizzled-3, a new *Drosophila* Wnt receptor, acting as an attenuator of Wngless signalling in *wingless* hypomorfic mutants. Development, 126, 4421–4430.

Shetty, P., Lo, M. C., Robertson, S. M., & Lin, R. (2005). *C. elegans* TCF protein, POP-1, converts from repressor to activator as a result of Wnt-induced lowering of nuclear levels. Developmental Biology, 285(2), 584–592. 10.1016/j.ydbio.2005.07.008

Sugioka, K., Mizumoto, K., & Sawa, H. (2011). Wnt regulates spindle asymmetry to generate asymmetric nuclear β-catenin in *C. elegans*. Cell, 146(6), 942–954. 10.1016/j.cell.2011.07.043

Sulston, J. E., & Horvitz, H. R. (1977). Post-embryonic cell lineages of the nematode, *Caenorhabditis elegans*. Developmental Biology, 56(1), 110–156. 10.1016/0012-1606(77)90158-0

Takeshita, H., & Sawa, H. (2005). Asymmetric cortical and nuclear localizations of WRM-1/β-catenin during asymmetric cell division in *C. elegans*. Genes and Development, 19(15), 1743–1748. 10.1101/gad.1322805

van der Horst, S. E. M., Cravo, J., Woollard, A., Teapal, J., & van den Heuvel, S. (2019). *C. elegans* Runx/CBFβ suppresses POP-1 TCF to convert asymmetric to proliferative division of stem cell-like seam cells. Development, 146(22), 1–14. 10.1242/dev.180034

Vora, S., & Phillips, B. T. (2015). Centrosome-associated degradation limits β-catenin inheritance by daughter cells after asymmetric division. Current Biology, 25(8), 1005–1016. 10.1016/j.cub.2015.02.020

Whitacre, J. M. (2012). Biological robustness: Paradigms, mechanisms, systems principles. In Frontiers in Genetics (Vol. 3, Number MAY). 10.3389/fgene.2012.00067

Wildwater, M., Sander, N., de Vreede, G., & van den Heuvel, S. (2011). Cell shape and Wnt signaling redundantly control the division axis of *C. elegans* epithelial stem cells. Development, 138(20), 4375–4385. 10.1242/dev.066431

Willert, J., Epping, M., Pollack, J. R., Brown, P. O., & Nusse, R. (2002). A transcriptional response to Wnt protein in human embryonic carcinoma cells. BMC Developmental Biology, 2.

Yang, X. D., Huang, S., Lo, M. C., Mizumoto, K., Sawa, H., Xu, W., Robertson, S., & Lin, R. (2011). Distinct and mutually inhibitory binding by two divergent β-catenins coordinates TCF levels and activity in *C. elegans*. Development, 138(19), 4255–4265. 10.1242/dev.069054

Yang, X. D., Karhadkar, T. R., Medina, J., Robertson, S. M., & Lin, R. (2015). β-Catenin-related protein WRM-1 is a multifunctional regulatory subunit of the LIT-1 MAPK complex. PNAS, 112(2), E137–E146. 10.1073/pnas.1416339112

